# Identifying determinants of synaptic specificity by integrating connectomes and transcriptomes

**DOI:** 10.1101/2023.04.03.534791

**Authors:** Juyoun Yoo, Mark Dombrovski, Parmis Mirshahidi, Aljoscha Nern, Samuel A. LoCascio, S. Lawrence Zipursky, Yerbol Z. Kurmangaliyev

**Author notes:** Corresponding authors. (S.L.Z); (Y.Z.K.). Department of Biology, Brandeis University; Waltham, MA 02453, USA.

## Abstract

How do developing neurons select their synaptic partners? To identify molecules matching synaptic partners, we integrated the synapse-level connectome of neural circuits with the developmental expression patterns and binding specificities of cell adhesion molecules (CAMs) on pre- and postsynaptic neurons. We focused on parallel synaptic pathways in the Drosophila visual system, in which closely related neurons form synapses onto closely related target neurons. We show that the choice of synaptic partners correlates with the matching expression of receptor-ligand pairs of Beat and Side proteins of the immunoglobulin superfamily (IgSF) CAMs. Genetic analysis demonstrates that these proteins determine the choice between alternative synaptic targets. Combining transcriptomes, connectomes, and protein interactome maps provides a framework to uncover the molecular logic of synaptic connectivity.

## Introduction

Advances in electron microscopy (EM) level connectomics have demonstrated the extraordinary complexity and specificity of synaptic connectivity ^1–7^. Developing neurons encounter the axons and dendrites of many different neuron types and form synapses with only a subset of them. Considerable progress has been made in understanding the molecular mechanisms by which neurons reach their target regions (axon guidance ^8^) and form synapses (synaptogenesis ^9^). Here we focus on the less well-understood step of synaptic specificity, how neurons choose their synaptic partners ^10^.

Sperry proposed that molecular labels allow neurites to discriminate between one another ^11^. Identifying these labels and understanding how they work is key to uncovering the molecular basis of brain wiring. Neurons express complex repertoires of many different cell surface molecules that mediate heterophilic and homophilic binding ^12–16^. These molecules, including members of the SYG/Kirre, Dscams, Cadherins, DIPs/Dprs and other families, contribute to synaptic specificity in different ways ^10,17–21^. It has been proven difficult, however, to identify matching receptor-ligand pairs expressed on pre- and postsynaptic membranes that lead to the discrimination between alternative synaptic partners and, thus, specify synaptic connectivity ^22^.

We address this challenge by combining the synaptic connectome of the neural circuits with the developmental transcriptome of the neurons comprising them. Combining these data provides a detailed description of all the CAMs expressed by developing pre and postsynaptic partners during the assembly of the connectome. These data can be further augmented by protein-protein interactomes to chart possible molecular interactions between CAMS expressed on developing synaptic partners. Typically, there are numerous potential interacting proteins on the surface of any two neuron types and, thus, identifying candidates as regulators of specificity in this way has proven difficult. To identify matching receptor-ligand pairs regulating synaptic specificity we focus on pairs of closely related neurons that choose different synaptic targets. Using this approach, we identified molecular determinants of synaptic specificity in the motion detection circuit of Drosophila melanogaster ^23^.

## Results

### Coupled transcriptome-connectome map of the Drosophila motion detection circuit

Dense EM reconstructions identified most of the synaptic connections in this circuit ^3,7^ (**Figures 1A** and **1B**). A set of eight closely related subtypes of T4/T5 neurons lie at its center. Each subtype is defined by a combination of one of two patterns of dendritic inputs and one of four patterns of axonal outputs (**Figures 1A** and **1F**). T4 and T5 neurons arborize their dendrites in the medulla and lobula, respectively. Each of these groups is further subdivided into four subtypes (a/b/c/d) based on their axon terminals in four synaptic layers of the lobula plate (Lop1/2/3/4). Each pair of T4 and T5 neurons which project axons to the same layer respond optimally to motion in one cardinal direction; T4 neurons respond to the movement of bright edges (ON pathway) and T5 neurons respond to dark edges (OFF pathway) ^24^. T4 and T5 axons terminating in the same layer converge onto the same postsynaptic partners ^7^. Furthermore, some of the postsynaptic neurons in different layers are also closely related cell types (see below). In this way, information from the ON (T4) and OFF (T5) pathways corresponding to each cardinal direction converge onto four parallel synaptic pathways (**Figure 1B**). We hypothesize that T4 and T5 subtypes, which form synapses with the same set of postsynaptic neurons in each layer of the lobula plate, do so through the same molecular mechanisms ^25^.

**Figure 1.**
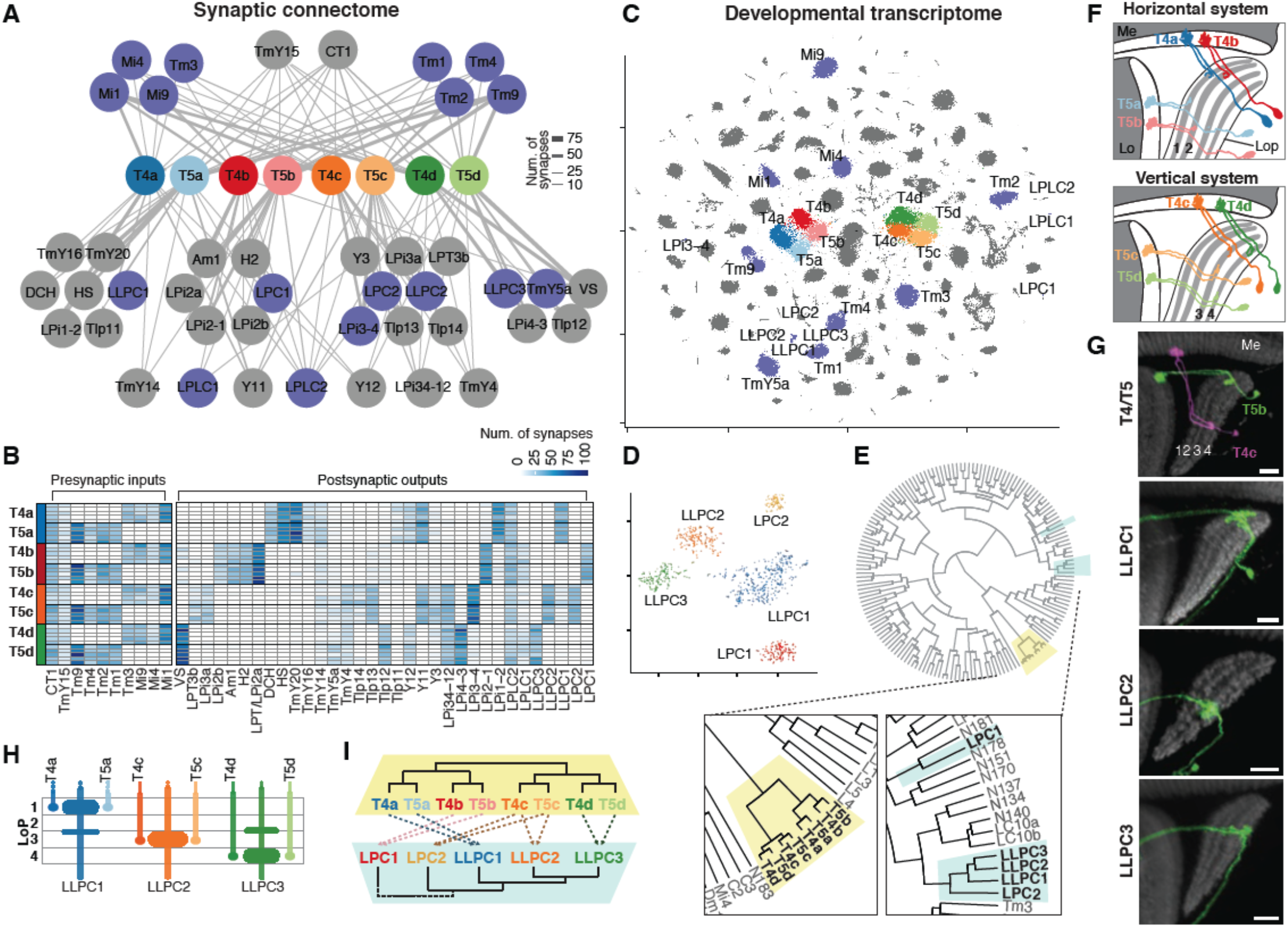
Coupled transcriptome-connectome map of the Drosophila motion detection circuit. (A-B) The connectome of T4/T5 subtypes from ref. ^3,7^. (A) Cell-type level connectome graph. Presynaptic inputs (top) and postsynaptic outputs (bottom) of T4/T5s are grouped and averaged by cell types. Purple, cell types with known transcriptomes. (B) Connectomes of individual T4/T5s, as an adjacency matrix. Five instances for each T4/T5 subtype (rows). Synaptic partners are grouped by cell type (columns). Each pair of T4 and T5 subtypes converge onto the same set of postsynaptic targets. (C) tSNE of the Drosophila visual system atlas ^14^. T4/T5s and their synaptic partners are labeled as in A. (D) LPC/LLPC clusters. See also Figure S1. (E) Hierarchical clustering of transcriptomes in the visual system atlas; T4/T5 and LPC/LLPC neurons are shown in the zoom-in. (F) Morphology of eight T4/T5 subtypes. Me, medulla; Lo, lobula; LoP, lobula plate. (G) Sparsely labeled T4/T5s and LLPC neurons. Neuropil marker (grey), brp. Scale bars, 10 μm. (H) Layer targeting of T4/T5 and LLPC neurons in the LoP. For LPCs see Figure S1 (I) Closely related pairs of T4 and T5 subtypes converge onto the same postsynaptic targets.

To identify these molecules, we integrated the synaptic connectome and the transcriptome of developing neurons in the Drosophila visual system (**Figures 1A-1C**). We previously generated a comprehensive transcriptional atlas of the developing visual system using single-cell RNA sequencing ^14^. This atlas covers more than 160 neuronal populations at seven developmental time points; ∼100 of them were matched to cell types in the connectome (ref ^26^; A.N., Y.Z.K, S.L.Z., *in preparation*). This includes transcriptional profiles of all T4/T5 subtypes and 17 of their synaptic partners (**Figure 1C**). We focus on five types of morphologically similar postsynaptic partners ^7,26^: two LPC (Lobula Plate Columnar) and three LLPC (Lobula-Lobula Plate Columnar) neurons **(Figures 1D** and **S1**). Each of these neuron types receives its major input from one pair of T4 and T5 subtypes (in one layer of the lobula plate) (**Figures 1B** and **1G**). In our initial version of the transcriptional atlas these five cell types were not resolved. A more detailed analysis revealed distinct transcriptional clusters for each of them, which were validated by *in vivo* expression patterns of marker genes (**Figures S1** and **S2**) and mRNA profiling of purified cell types (A.N., Y.Z.K, S.L.Z., *in preparation*). Hierarchical clustering of transcriptomes of all neuronal populations in the visual system confirmed that the eight T4/T5 subtypes were closely related. Similarly, four LPC/LLPC types (except LPC1) were also closely related to each other (**Figure 1E**). Taken together, T4/T5 subtypes and LPC/LLPC types assemble into parallel synaptic pathways comprising homologous pairs of pre- and postsynaptic partners (**Figures 1H** and **1I**).

### Beat/Side IgSF CAM expression defines synaptic specificity

To understand how neurons choose their synaptic partners, we focused on synapses between the most closely related T4/T5 pairs and their closely related postsynaptic targets. T4c and T5c form synapses with LLPC2, and T4d and T5d form synapses with LLPC3 (**Figure 2A**). Each of these pairs of T4/T5 subtypes converge onto transcriptional programs that correlate with the specificity of their axonal outputs ^25^. Virtually all differentially expressed genes (DEGs) between T4c and T4d are the same as between T5c and T5d (**Figure 2B**). There were only nine such DEGs, and of these, seven were cell surface proteins (**Figure 2C**). Similarly, a small number of cell surface proteins were differentially expressed between LLPC2 and LLPC3 neurons (**Figure 2D**). We hypothesize that matched pairs of cell recognition molecules specific to each pair of synaptic partners regulates this binary choice in synaptic specificity.

**Figure 2.**
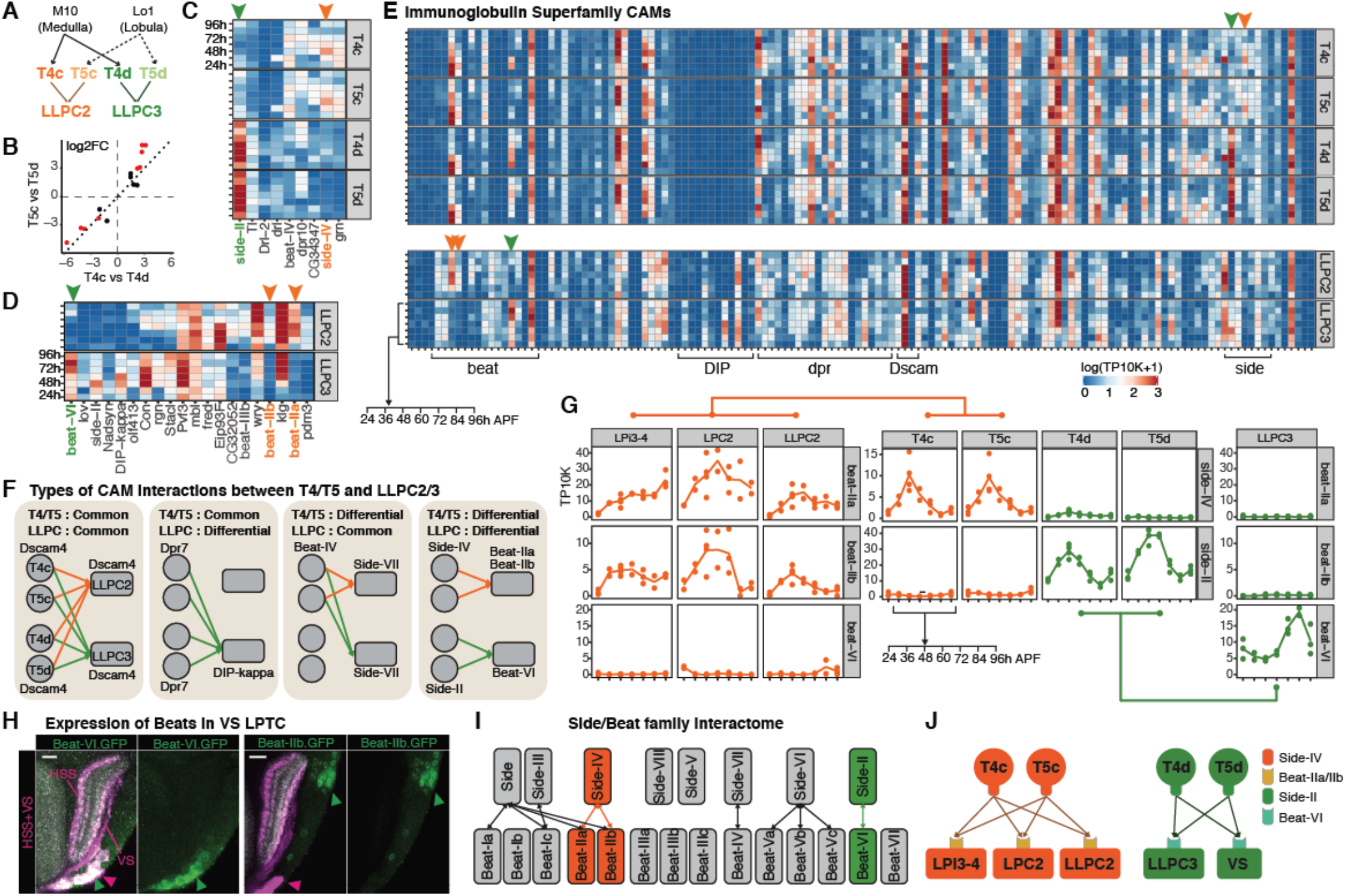
Matching receptor-ligand pairs correlate with synaptic specificity. (A) Connectome of T4/T5-LLPC2/3 circuit. (B to D) Differentially expressed genes (DEGs) between T4/T5 and LLPC subtypes. See Methods for details and thresholds. (B) Log-fold-change for DEGs between T4c and T4d (x-axis), and T5c and T5d (y-axis). (C) Heatmap of expression patterns of DEGs significant in both comparisons in B (red). (D) DEGs between LLPC2 and LLPC3 (E) All annotated IgSF CAMs. Data is shown for 7 time points after pupal formation (APF). (F) Examples of matching CAM interactions between T4/T5 and LLPC. Only two pairs of interactors correlate with connectome: Side-IV::Beat-IIa/b and Side-II::Beat-VI (orange and green arrows in C to E). (G) Line plots with expression patterns of these Beat/Side proteins in the T4/T5-LLPC2/3 circuit, and other major targets with known transcriptomes. Dots are replicates. (H) *In vivo* expression of Beat-IIb and Beat-VI in VS neurons (adult), the main target of T4d/T5d (Figure 1D). Neuropil marker (grey), brp. Scale bars, 10 μm. (I) Interactome of Beat/Side families from ref. ^13^. (J) Matching receptor-ligand pairs between T4/T5s and their targets in Lop3 (orange) and Lop4 (green).

Although each T4/T5 subtype and LLPC types express many CAMs (**Figure 2E**), only two pairs of interacting CAMs correlated with synaptic specificities of these two sets of synaptic partners (**Figure 2F**). Both pairs belong to the Side and Beat families of IgSF proteins, which form a heterophilic protein interaction network ^13,27^ (**Figure 2I**). Founding members of these families (Side and Beat-Ia) were identified in genetic screens as regulators of motor axon guidance in the Drosophila embryo ^28–30^. Functions for other paralogs have not been described. As each neuron type in the developing visual system expresses a unique combination of 14 Beat and eight Side proteins during development, these proteins may contribute to synaptic specificity more broadly (**Figure S3**).

The top differentially expressed genes between T4/T5 subtypes are side-IV (specific to T4c/T5c) and side-II (specific to T4d/T5d). LLPC2 and LLPC3 neurons express interacting Beats in a matching fashion; beat-IIa and beat-IIb are specific to LLPC2, and beat-VI is specific to LLPC3 (**Figure 2G**). At least two other major synaptic targets of T4c/T5c (LPC2 and LPi3-4) express beat-IIa/IIb but not beat-VI, and the main synaptic target of T4d/T5d (VS) expresses Beat-VI but not Beat-IIb (**Figures 2G** and **2H**). In this way, the matching expression of two pairs of interacting IgSF CAMs correlates with synaptic specificity in this circuitry (**Figure 2J**).

### Side-II/Beat-VI is required for synaptic layer separation

We sought to assess the roles of Beat/Side interactions in the wiring of T4/T5 axons. In wild-type, these axon terminals form four layers in the lobula plate (**Figure 3A**). In homozygous *side-II*^null^ mutant animals, T4/T5 axon terminals formed a single fused Lop3/4 layer (**Figure 3B**). Lop1 and Lop2 were normal. Beat-VI is a high-affinity binding partner of Side-II. Homozygous *beat-VI*^null^ mutants phenocopy *side-II*^null^ mutants. These results were confirmed with an insertion and a deletion disrupting these genes (Figures S4A and S4B). The layer fusion phenotype resembles the earlier stages of development (**Figure 3A**).

**Figure 3.**
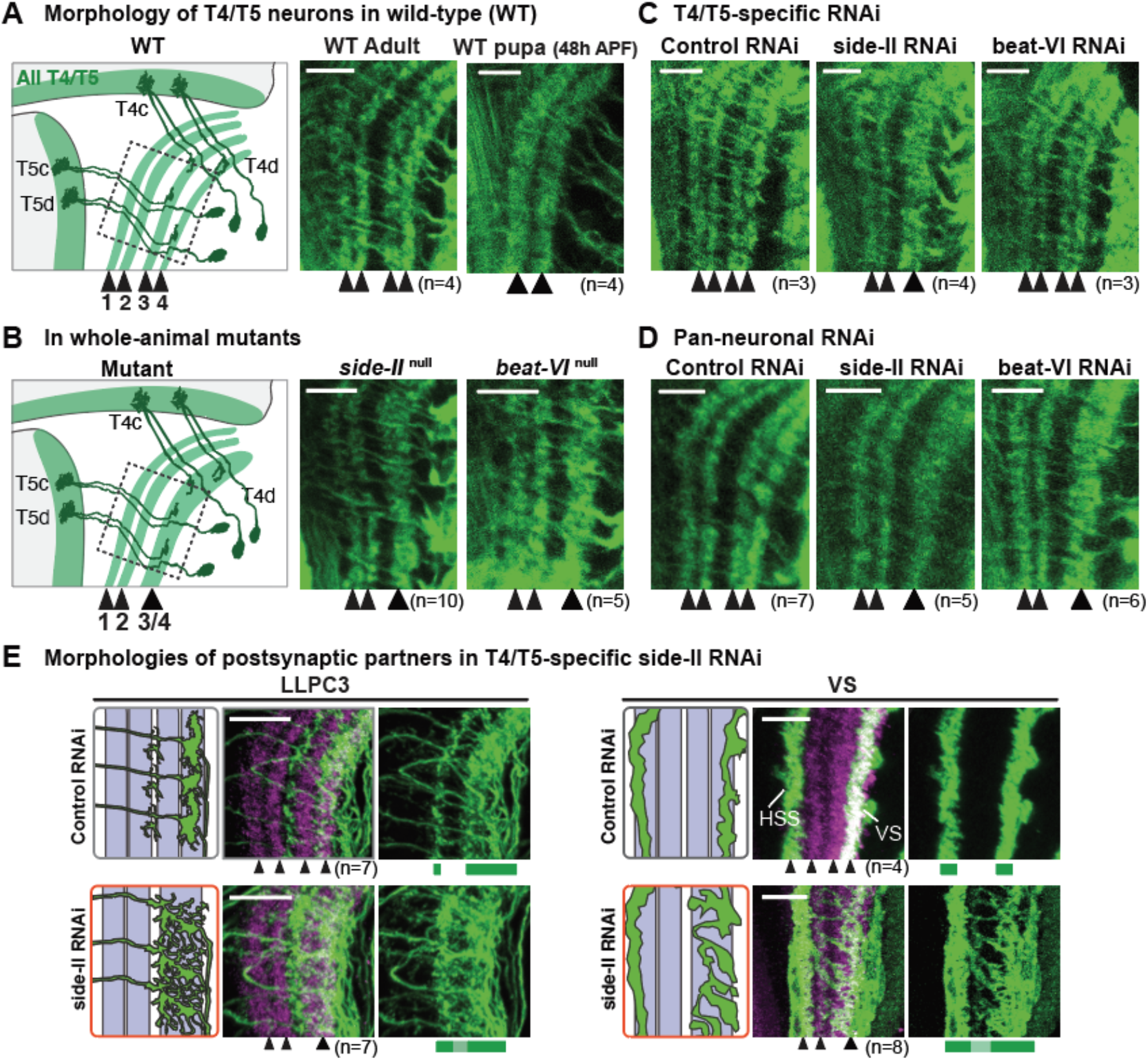
Side-II::Beat-VI segregates synaptic partners into layers. (A to D) Morphology of T4/T5s in the wild-type and mutant backgrounds. In immunofluorescence images, all T4/T5 neurons are labeled. (A) During development (*right*), axon terminals first laminate into two broad domains (Lop1/2 and Lop3/4). In adults (*middle*), these layers further separate into four layers (Lop1/2/3/4). (B) In *side-II*^null^ and *beat-VI*^null^ mutants, axons of T4c/T5c and T4d/T5d form a single fused Lop3/4 layer. (C) T4/T5-specific RNAi of Side-II (but not Beat-VI) also results in fused Lop3/4. (D) Pan-neuronal RNAi of both Side-II and Beat-VI results in fused Lop3/4. (E) Morphology of the main postsynaptic partners of T4/T5s (green) upon removal of Side-II from T4/T5s (via RNAi). In controls, LLPC3 and VS arborize in Lop4. In side-II RNAi, they span fused Lop3/4 layer. Neuropil marker (magenta), brp. Scale bars, 10 μm.

Next, we tested whether Side-II was required in T4/T5 neurons using RNA interference (RNAi) (**Figures 3C** and **3D**). Removing side-II specifically from T4/T5s phenocopied *side-II*^null^ mutants. These results were confirmed using an independent side-II RNAi line (**Figure S4A**). Removing beat-VI from T4/T5s did not result in the fusion of Lop3/4, whereas a pan-neuronal RNAi of beat-VI did. Removal of side-II from T4/T5s also disrupted dendritic morphologies of their main postsynaptic partners (**Figure 3E**). Thus, Side-II and Beat-VI is a receptor-ligand pair that regulates separation of neuronal processes into adjacent synaptic layers of the lobula plate.

Expression patterns of Side-IV and Beat-IIa/IIb in Lop3 suggested a similar role for this receptor-ligand pair in lobula plate development. However, in homozygous *side-IV*^null^ mutant animals, we did not observe defects in the lamination of T4/T5 axon terminals (**Figure S4C**). This suggests that these proteins have a different function, or other recognition molecules may act to compensate for their loss.

### Side-II and Beat-VI control synaptic specificity

Next, we assessed the role of Side-II/Beat-VI interactions at the level of single cells. For T4/T5 neurons, we visualized the morphologies and presynaptic sites of sparsely distributed homozygous *side-II*^null^ mutant neurons where most, if not all, other neurons are wild-type (heterozygous; see Methods, **Figures 4A-4C**). Wild-type T4c/T5c and T4d/T5d have presynaptic sites in Lop3 and Lop4, respectively. Mutant T4d/T5d axons terminate in Lop4, however, their presynaptic sites span both layers. The penetrance of this phenotype was complete (17 of 17 single mutant neurons). Mutant T4c/T5c were similar to wild-type. The simplest interpretation is that mutant T4d/T5d neurons form ectopic synapses with inappropriate synaptic partners in Lop3.

**Figure 4.**
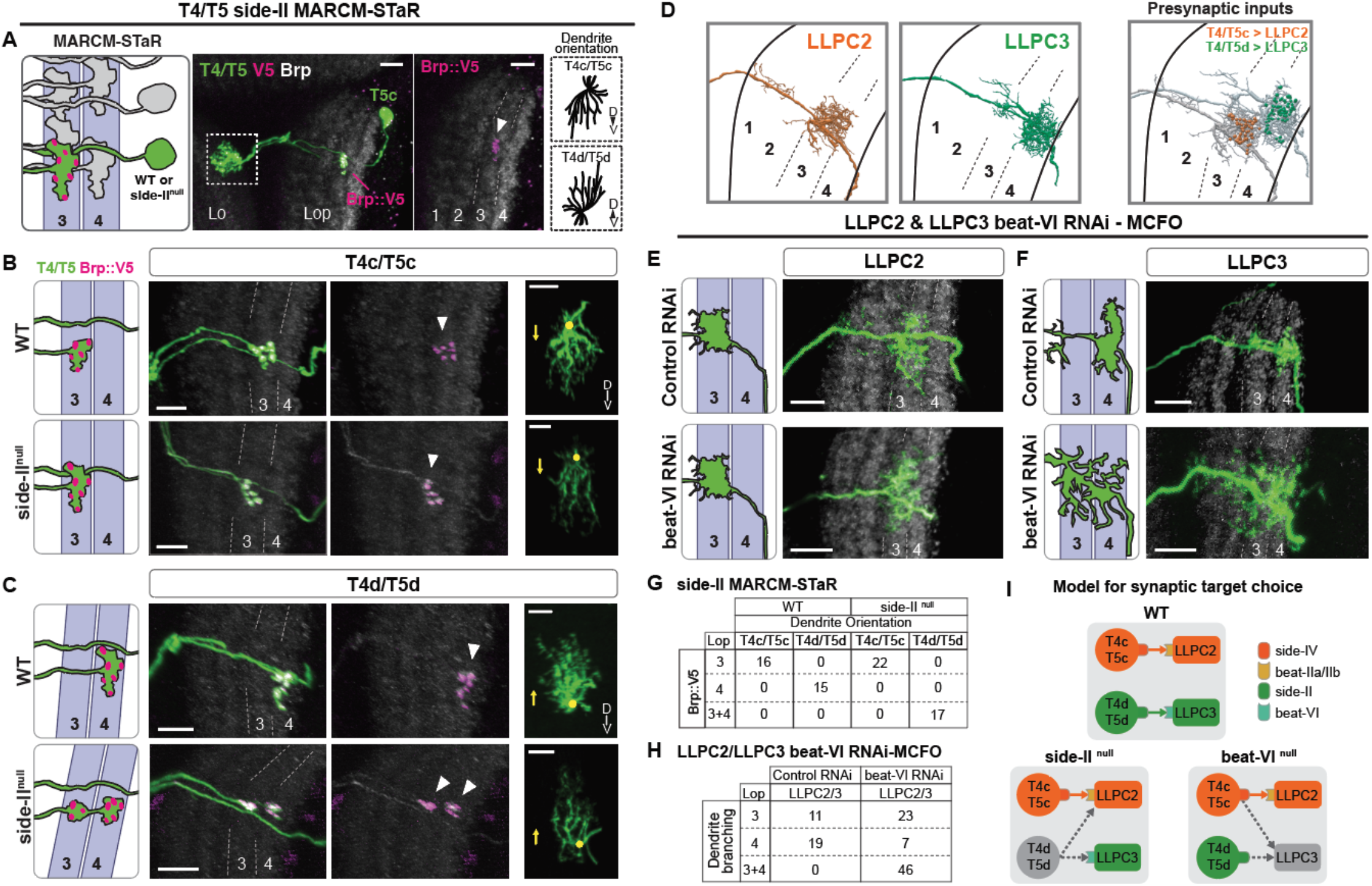
Side-II::Beat-VI determines the choice between alternative synaptic targets. (A to C) MARCM-STaR. Mosaics with homozygous mutant (*side-II*^null^) T4/T5 neurons (green) with presynaptic marker Brp-V5 (magenta), and wild-type controls (WT). (left) Schematics of T4/T5 axon terminals in Lop3 and Lop4. (right) Dendrite orientation discriminates between T4c/T5c and T4d/T5d (D: dorsal, V: ventral, based on the visual field coordinates). (B) WT and *side-II*^null^ T4c/T5c form synapses in Lop3. (C) WT T4d/T5d form synapses in Lop4; *side-II*^null^ T4d/T5d form synapses in Lop3 and Lop4. (D) LLPC2 and LLPC3 morphologies and T4/T5 inputs from EM reconstruction ^7^. (E and F) beat-VI RNAi in LLPC2 and LLPC3. Single neurons were visualized using MCFO. In controls, both LLPC2 and LLPC3 dendrites were wild-type (as in D). In beat-VI RNAi, we detected wild-type LLPC2 and fewer LLPC3 than expected. We observed a large number of abnormal neurons spanning both Lop3 and Lop4. As beat-VI is specific to LLPC3, we conclude that abnormal neurons are LLPC3. (G) Quantification of the labeled neurons from B and C. (H) Quantifications of labeled neurons from E and F. (I) Model for circuit rewiring upon removal of Side-II and Beat-VI. Neuropil marker (gray), brp. Scale bars, 10 μm.

We next removed Beat-VI from postsynaptic partners, LLPC2 and LLPC3, using RNAi and visualized individual mutant neurons (see Methods, **Figures 4E-4G**). In controls, we detected wild-type LLPC2 (dendrites in Lop3) and LLPC3 (dendrites in Lop4). In beat-VI RNAi, we detected wild-type LLPC2 and many abnormal neurons spanning both Lop3 and Lop4. Fewer wild-type LLPC3s than expected were observed (**Figure 4H**, Fisher’s exact test, p = 0.004). As beat-VI is specific to LLPC3, we conclude that abnormal neurons are mutant LLPC3s. This phenotype was confirmed using an independent beat-VI RNAi line (**Figure S5**). The single-cell phenotypes are consistent with each other. That is, presynaptic sites of Side-II mutant T4d/T5d and postsynaptic dendrites of Beat-VI-deficient LLPC3 are no longer restricted to Lop4 and accumulate in Lop3.

In summary, Side-II is expressed in T4d/T5d axons and Beat-VI in dendrites of their postsynaptic partners. Side-II and Beat-VI bind to each other. Removal of this receptor-ligand pair leads to defects in synaptic circuitry in Lop4. These proteins may control several steps of circuit assembly. At an early stage, they promote adhesion between axons and dendrites leading to their segregation into layers. At a later time, they may directly specify connections between synaptic partners. Regardless of mechanistic details, Side-II and Beat-VI interactions play a critical role in matching synaptic partners.

## Discussion

How do neurons choose their synaptic partners? We have shown that any pair of neurons typically express dozens of CAMs that can promote molecular interactions between them ^14^. Removing single genes frequently leads to only modest defects in connectivity^31^. This has led to the view that selective wiring between pre- and postsynaptic neurons requires many genes each contributing in only a small way to specificity ^31^. Alternatively, for instance, this may reflect the difficulty in assessing defects in synaptic specificity in the densely packed CNS. Our data support this latter view. Defects in synaptic specificity of side-II mutants were only uncovered by visualizing the morphologies and presynaptic sites of sparsely distributed single mutant T4d/T5d neurons. The single synaptic phenotypes exhibited complete penetrance. That these neurons also form synapses in their normal target Lop4, however, underscore the contributions of other pathways, and thus redundancy, to regulating synaptic specificity.

Here we demonstrate that specificity can be determined by a differential match in a single receptor-ligand pair. If we remove these interactions the molecular differences between correct and incorrect targets diminish and they become less distinct. For instance, in the absence of Side-II, LLPC2 and LLPC3 would appear more similar to T4d/T5d than in wild-type (**Figure 4C**). As a consequence, T4d/T5d forms synapses in Lop3, in addition to Lop4. During the circuit assembly neurons must discriminate between many potential synaptic partners in a neuropil comprising densely intermingled processes of many different cell types. Our findings support the idea that the interactions between the vast number of CAMs expressed on the surfaces of neurons control a hierarchy of wiring decisions sequentially restricting the pool of possible targets ^32,33^, The last step in this process is the selection of synaptic partners and this is determined by the relative preference between potential synaptic partners ^34^.

The work we describe here expands the diverse repertoire of families of IgSF proteins which contribute to synaptic specificity in the *Drosophila* brain ^17,21^. Each family forms complex receptor-ligand networks including homophilic (e.g. Dscams ^35^ thousands of isoforms) and heterophilic interactions (e.g. DIP/Dpr ^27,36^ and Side/Beat ^13,27,30^ comprising 50+ interacting pairs). These proteins are expressed in highly dynamic and cell-type-specific ways, and with other cell surface proteins endow each neuron with a unique cell surface protein composition ^11,14^. The logic in wiring the mammalian brain may be similar, with an expanded cadherin superfamily largely taking the place of IgSF diversity ^10^.

Coupled transcriptome-connectome maps provide a description of gene expression patterns for both sides of synaptic connections. These maps can be correlated with binding specificities of cell surface proteins to chart possible molecular interactions between neurons ^37,38^. As more connectomes ^4,39^ and developmental transcriptomes ^40^ become available, comparative studies of highly related groups of neurons with divergent wiring specificities may prove fruitful in uncovering matched receptor-ligand pairs regulating synaptic specificity in the mammalian brain.

## Methods

### Connectome analysis

T4/T5 connectomes ^3,7^ were downloaded from the neuPrint database (https://neuprint.janelia.org/, dataset: fib19:V1.0, ^41^) using natverse (0.2.4) package ^42^. Synaptic connectivity data (inputs and outputs) were downloaded for 40 representative T4/T5 neurons from published data ^7^ (five instances of each T4/T5 subtype). For each representative neuron, we summed the total number of synapses between a given instance of T4/T5 and all synaptic partners of the same cell type (e.g. total number of synapses between a single T4a and any Mi1 neurons). This data was plotted as a heatmap of synaptic weights between partner neuron types and individual instances of T4/T5 neurons (**Figure 1B**). Synaptic weights were averaged across all instances of each T4/T5 subtype to generate a cell type-level connectome graph. The connectome graph was visualized using Cytoscape ^43^ and igraph ^44^ (**Figure 1A**). Synaptic partners were restricted to cell types that make more than 10 synapses with a single T4/T5 neuron. Only connections with more than 7 synapses have been plotted. Presynaptic inputs were restricted to connections in the medulla and lobula; postsynaptic outputs were restricted to connections in the lobula plate. Reconstructions of representative LLPC2 and LLPC3 neurons in Fig. 4D were visualized in neuPrint.

### Transcriptome analysis

#### Analysis of the transcriptional atlas of the Drosophila visual system

Single-cell analysis was performed using Seurat (V4.1.1 ^45^). All functions were used with default parameters, unless otherwise indicated. In this study, we use single-cell RNA-Seq data from our previously generated comprehensive transcriptional atlas of the developing visual system ^14^. We focus on the main dataset including samples from seven developmental time points taken every 12 hours from 24 to 96h APF. In the initial version of the visual system atlas (V1.0), 58 transcriptional clusters were matched to known morphological cell types. Two of these clusters were annotated as LLPC1 and LPC1 neurons based on correlation analysis with available bulk reference datasets ^26^. A more detailed evaluation of these clusters revealed further heterogeneity in the LLPC1 cluster. We subsetted and reclustered LLPC1 and LPC1 clusters separately from the rest of the dataset. This analysis was performed after the integration of the main dataset as previously described ^14^. After subsetting, a new set of 1000 highly variable genes was selected, scaled, and used for principal component analysis (functions: FindVariableFeatures, ScaleData, RunPCA). The first nine principal components were used for the generation of tSNE plots and clustering (functions: RunTSNE, FindNeighbors, FindClusters, resolution = 0.1). The analysis revealed five transcriptionally distinct clusters of LPC/LLPC neurons. Clusters were annotated based on the expression of cell-type-specific marker genes (**Figure S2**). Cell types of LPC/LLPC neurons were renamed in the main dataset of the visual system atlas (V1.1).

#### Visualization of gene expression patterns

Expression patterns of genes were visualized using average expression levels for each cell type and time point. Averaging was performed for each replicate in non-log space for the original normalized expression values (TP10K, transcripts-per-10,000). For the heatmaps, we used log1p-transformed expression values averaged across replicates (capped at the maximum expression value of 20). For the line plots, we used expression values in linear scale. The list of all IgSF proteins was obtained from FlyXCDB (http://prodata.swmed.edu/FlyXCDB;^46^).

#### Hierarchical clustering of neuronal cell types

Clustering was performed for neuronal cell types from one sample at 48h APF (W1118 sample, replicate B). This analysis was performed based on the original normalized expression values (pre-integration). We selected 1000 highly variable genes and computed average expression levels for each cell type (functions: FindVariableFeatures, AverageExpression). Hierarchical clustering was performed based on Pearson’s correlation coefficients between log-transformed expression profiles of each cell type (distance metric: 1 - Pearson’s r; clustering method: ward.D2) ^47^. Clustering results were visualized as a dendrogram (function: ape::plot.phylo) ^48^.

#### Differential gene expression analysis

We identified differentially expressed genes using Wilcoxon rank-sum test (function: FindMarkers, min.pct = 0.35, pseudocount.use = 0.01, max.cells.per.ident = 1000, fold-change > 2, adjusted p < 0.01). Marker genes common to all LPC/LLPC neurons were identified by comparison of all LPC/LLPC clusters to all other neurons in the atlas; cell-type-specific markers were identified by comparison of individual LPC/LLPC clusters to other LPC/LLPC neurons. Marker genes were identified for all time points and replicates together. Expression patterns of select markers are shown in Figure S2. Differential analysis of T4/T5 and LLPC neurons in Lop3 and Lop4 for Fig. 2 was performed at 48h APF (pseudocount.use = 0.01, fold-change > 3, adjusted p < 0.01).

The analysis of connectome and transcriptome data was performed in R (4.1.0). Single-cell RNA-Seq data used in this study and new cell type annotations for the visual system atlas (V1.1) are available through NCBI GEO: GSE156455. The code used in this work is available on request.

### Fly Genetics

Flies were kept on standard cornmeal medium at 25°C, 12-hour light/dark cycles. All RNAi experiments were run with UAS-Dcr2.D. Detailed genotypes for all figures and supplementary material are included in a separate spreadsheet (Table S1)

#### Generation of null alleles using CRISPR

For side-II and beat-VI, two protospacer sequences targeting the first coding exon were chosen to create a short deletion leading to a frameshift mutation of the protein sequence. For side-IV, two protospacer sequences spanning the whole gene were chosen to create a total 12.9kb deletion. High score protospacer sequence was chosen from UCSC Genome Browser crisprTarget table. Oligos were made from selected gRNA sequences and inserted into pU6-2 vector ^49^. gRNA plasmid was injected into vas-Cas9 line (BDSC 51323) via Bestgene Inc. Injected larvae were crossed with balancer lines and F1 progeny was screened for mutation. A mutant allele line was established from this single F1 progeny. sgRNA sequences are listed. Detailed protocols are available upon request. side-II[13] (side-II null) deleted sequence(44bp): TCCGGCGGAGGCAGCAGCATGGGTCCTGGCGGAGGAGGATCCGG side-IV[4-5] deleted sequence(12.9kb): AACGCGTATTCGCACCCACACACAAGTGAAGTCGGCTCT….…………….. GGAACTCTCCGGCACTCCGGTATTCCGGAATTCCGTTGCTCCGGTGGTC beat-VI[4] (beat-VI null) deleted sequence(19bp): AAGGATACGGAGCCGGCCA

#### MARCM-STaR experiment

MARCM-STaR labels cell morphology and the presynaptic active zone protein (Brp) in single homozygous null mutant neurons in otherwise heterozygous backgrounds ^50^. The *side-II*[13] allele was recombined with FRT40 for MARCM ^51^. Mitotic recombination was induced at the third instar larva stage with 37°C heat shock for 2-3 min. FLP activated the FRT-flanked stop signal resulting in expression of R recombinase under GAL4 control. R recombinase then excised sequences encoding the stop codon flanked by R-specific-recombination sites (RSRT). This resulted in the insertion of the V5 tag into Brp and due to the insertion of the following T2A site expression of the linked LexA coding sequence. LexA then induces expression from LexAop-myr-tdTOM to label the cell membranes to highlight cell morphology. (Figure. S6, detailed genotype in Table S1). Flies were dissected within two days after eclosion. The brains were visualized by immunofluorescence staining as described below.

The T4/T5 GAL4 driver marks all subtypes. In order to classify the identities of T4/T5 MARCM clones, we used the unique dendritic orientation of T4/T5s (i.e. T4c/T5c, extending dorsal to ventral; T4d/T5d, extending ventral to dorsal). Brains were mounted to have confocal image stacks along the dorsal to ventral axis, so that the Z-axis in the final volume corresponds to the D-V axis of the compound eye (i.e. the visual field). Images were analyzed in IMARIS to enable 3D visualization. For each T4/T5 dendrite, orientation was determined by the angle of the primary dendritic branch extending away from the axon shaft (i.e. extension away from the axon) and the position of distal tips of the dendrite. T4c/T5c and T4d/T5d dendrites were oriented in opposite directions.

#### RNAi-MCFO experiment

RNAi was expressed by a GAL4 driver expressed in both LLPC2 and LLPC3. To visual single LLPC2 and LLPC3 neurons we combined MultiColor FlipOut (MCFO) with beat VI knock down (ref.^52^ **Figures 4E and 4H**). Heat shock (8-10 min) was induced in the mid-pupal stage for sparse labeling.

#### Immunohistochemistry / Immunofluorescence and confocal microscopy

Brains were dissected in ice-cold Schneider’s Drosophila Medium (GIBCO #21720-024), and fixed in PBS containing 4% paraformaldehyde (PFA) for 25 min at room temperature (RT). Brains were washed three times with PBST (PBS containing 0.5% Triton X-100), and incubated in blocking solution (PBST containing 10% Normal Goat Serum) for at least 2 hr at RT prior to incubation with antibody. Brains were incubated sequentially with primary and secondary antibodies diluted in blocking solution for 2 days at 4°C, with 3 PBST washes followed by 2 hr incubation at RT in between and afterward. Then brains were mounted with Everbrite mounting media (Biotium #23001) or processed for DPX mounting (see below).

#### Antibody information

Primary antibodies and dilutions used in this study: chicken anti-GFP (1:1000, Abcam #13970), rabbit anti-dsRed (1:200, Clontech#632496), mouse anti-Nc82 (1:40, Developmental Studies Hybridoma Bank (DSHB) Nc82), chicken anti-V5 (1:300, Fortis Life Sciences #A190-118A), rabbit anti-HA (1:200, Cell Signaling Technology #3724), guinea pig anti-Pdm3 (1:20, a gift from John Carlson), and mouse anti-Br (1:20, Developmental Studies Hybridoma Bank (DSHB) 25E9.D7). Secondary antibodies and dilutions used in this study: goat anti-chicken Alexa Fluor 488 (AF488) (1:1000, Invitrogen #A11039), goat anti-rabbit AF568 (Invitrogen #A11011, 1:200), goat anti-guinea pig 568 (1:500, ThermoFisher #A11075), goat anti-mouse AF568 (1:500, ThermoFisher #A11031), and goat anti-mouse AF647 (1:500, ThermoFisher #A21235).

#### Tissue clearing and DPX mounting

Antibody stained brains were mounted with DPX following Janelia FlyLight protocol (https://www.janelia.org/project-team/flylight/protocols). Briefly, after secondary antibody wash as described above, brains were post-fixed with 4% PFA for at least 3 hr in RT. Brains were washed with PBS and mounted on polylysine-L coated coverslip and sequentially dehydrated in increasing concentration of ethanol (50%, 75%, 90%, 100%, 100%, 100%, 10-min each). Dehydrated brains on coverslip were incubated in Xylene (Fisher Scientific, X5-500) (5min x 3) for tissue clearing. Then the coverslip was embedded in DPX (Electron Microscopy Sciences, #13510) and cured in the chemical hood for more than 2 days before imaging. DPX mounting images are noted in the genotype table (Table S1)

#### Confocal microscopy

Immunofluorescence images were acquired using Zeiss LSM 880 confocal microscope with Zen digital imaging software. Optical sections or maximum intensity projections were level-adjusted, cropped and exported for presentation using ImageJ software (Fiji) or IMARIS 9 (Oxford Instruments). Reported expression patterns were reproducible across three or more biological replicates.

## Data and materials availability

Connectome and transcriptome datasets used in this study are available through Neuprint (neuprint.janelia.org) and NCBI GEO (GSE156455). Newly generated mutant alleles and split-Gal4 lines are available upon request.

## Acknowledgments

We thank Kazunori Shinomiya for sharing connectome data prior to publication. Stocks obtained from the Bloomington Drosophila Stock Center (NIH P40OD018537) and Vienna Drosophila Resource Center were used in this study. We also thank G. M. Rubin, H. J. Bellen, and Qi Xiao for the reagents and the Janelia FlyLight Project Team for some images. We thank members of the Zipursky lab for the critical discussion of the manuscript.

## Authors contributions

J.Y., A.N., S.L.Z and Y.Z.K., conceptualization; J.Y., M.D., P.M, A.N., S.A.L, Y.Z.K., investigation and formal analysis; J.Y., S.L.Z and Y.Z.K., writing.

---

**Figure S1.**
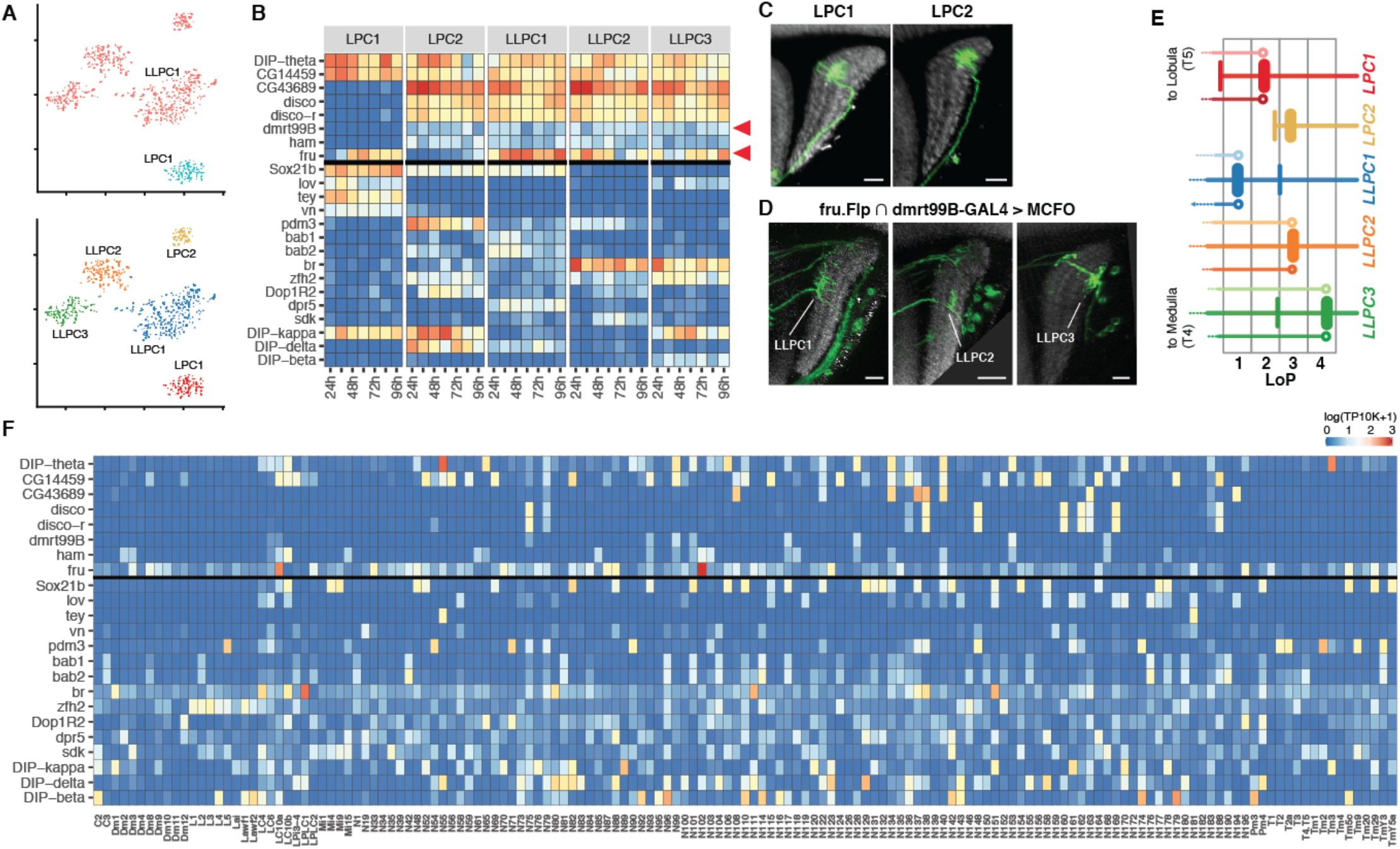
Transcriptomes of LPC/LLPC neurons. (A) tSNE plots of LPC/LLPC neurons. Cells are color-coded according to annotations in the atlas V1.0 (top) and V1.1 (bottom). In the first version of the atlas, all LLPC and LPC2 neurons were clustered together (as LLPC1). (B) Expression patterns of common and cell-type-specific markers of LPC/LLPC neurons. (C) Sparsely visualized LPC1 and LPC2 neurons. (D) The intersection of transcription factors *fru* and *dmrt99B* visualized by MCFO. Expression patterns of these genes are shown by arrows in B. As predicted, this intersection captures LLPC neurons. (E) Layer targeting of T4/T5 and LPC/LLPC neurons in the LoP. (F) Expression patterns of marker genes from C in all other neuronal clusters in the visual system atlas ^14^. Neuropil marker (gray), brp. Scale bars, 10 μm.

**Figure S2.**
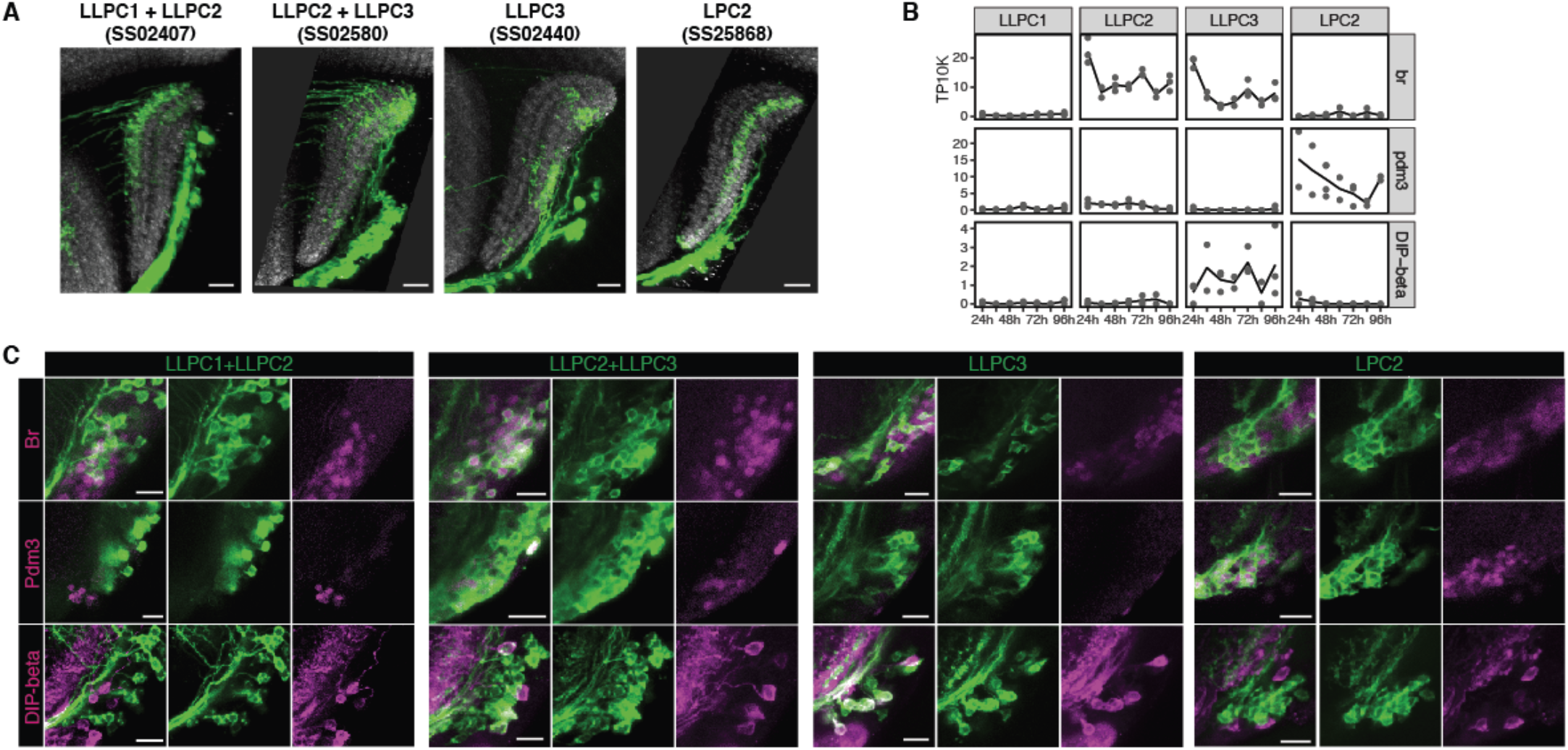
*In vivo* expression patterns of cell-type-specific marker genes of LPC/LLPC neurons. (A) Expression patterns of Gal4 drivers targeting subsets of LPC/LLPC types (green). (B) Expression patterns of select cell-type-specific marker genes in transcriptomes. (C) *In vivo* expression of marker genes from B (magenta), colocalized with LPC/LLPC cell bodies visualized using Gal4 drivers from A (green). Expression of *br* and *pdm3* is visualized using immunostaining for these proteins. Expression of DIP-beta is visualized using DIP-beta-lexA. Neuropil marker (gray), brp. Scale bars, 10 μm.

**Figure S3.**
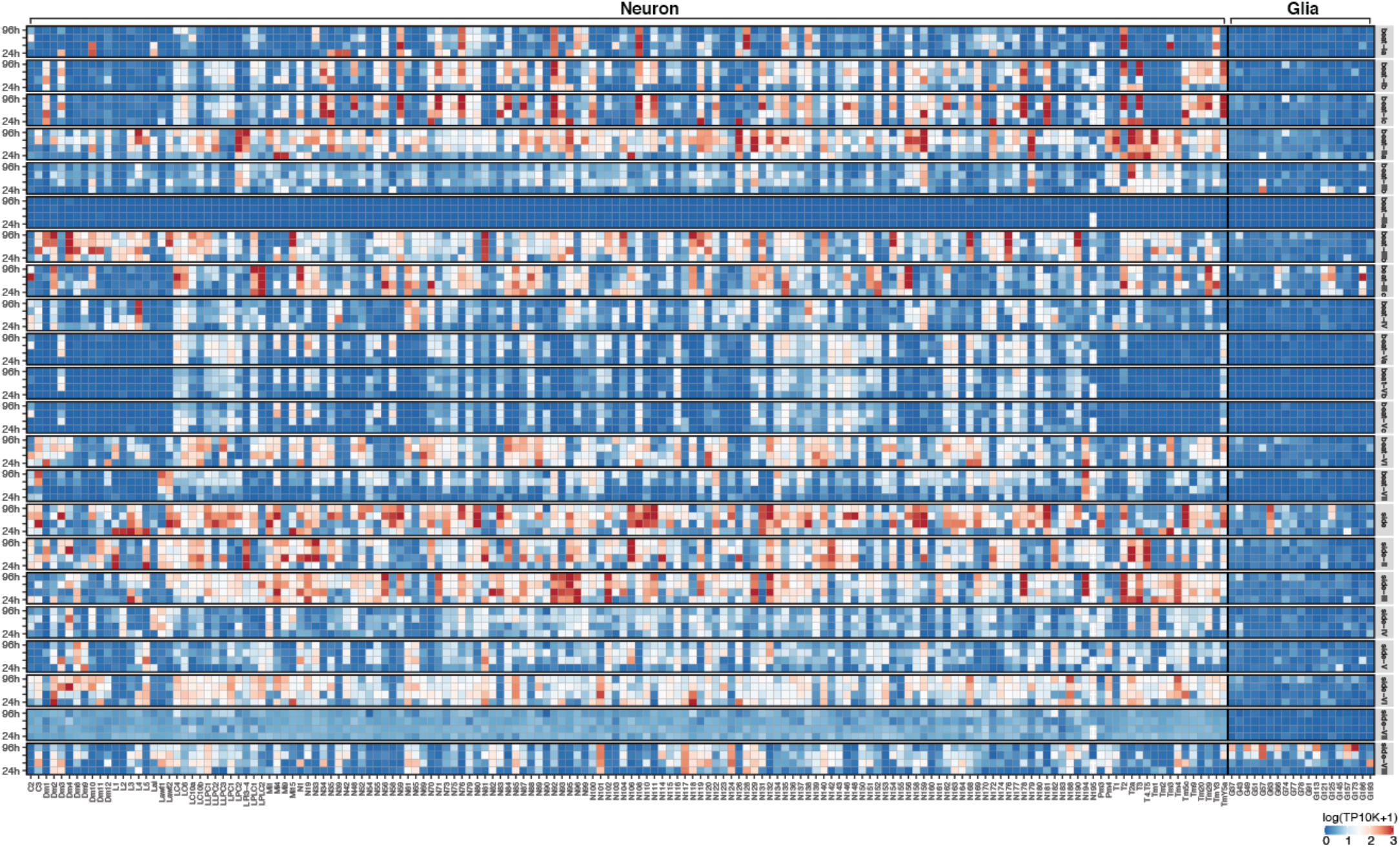
Expression patterns of Beat/Side families of proteins in the Drosophila visual system. Expression patterns are shown for all neuronal (left) and glial (right) clusters from the Drosophila visual system atlas ^14^. Data is shown for 4 time points (24h, 48h, 72h, and 96h APF). Lack of beat-IIIa expression could be due to technical reasons (e.g. misannotation of transcripts) or it is not expressed in the brain.

**Figure S4.**
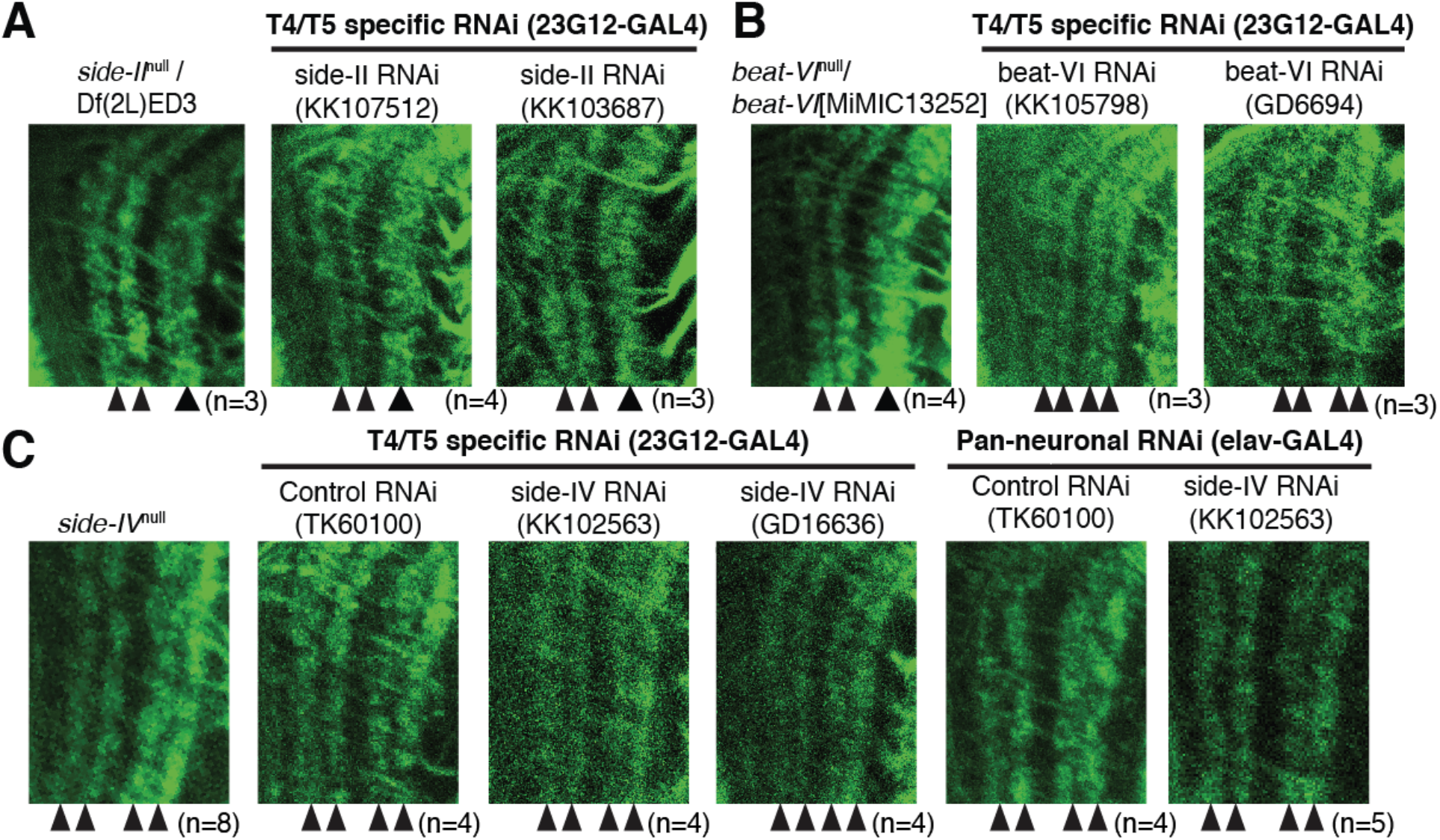
Side/Beat mutants. (A to C) Morphology of T4/T5 neurons in various mutant backgrounds, visualized as in Figure. 3. (A) *side-II*^null^ allele over deficiency and two independent side-II RNAi are indistinguishable from homozygous *side-II*^null^ animals (KK107512 is also shown in Figure 3). (B) *beat-VI*^null^ allele over a MiMIC insertion into a beat-VI gene phenocopies homozygous *beat-VI*^null^ animals. Two independent beat-VI RNAi lines expressed in T4/T5s are indistinguishable from controls. By contrast, the same RNAi lines expressed in all neurons phenocopies homozygous *beat-VI*^null^ animals (KK105798 is also shown in Figure 3). (C) In *side-IV*^null^ and side-IV RNAi, T4/T5 axons form four layers in the LoP as in wild-type.

**Figure S5.**
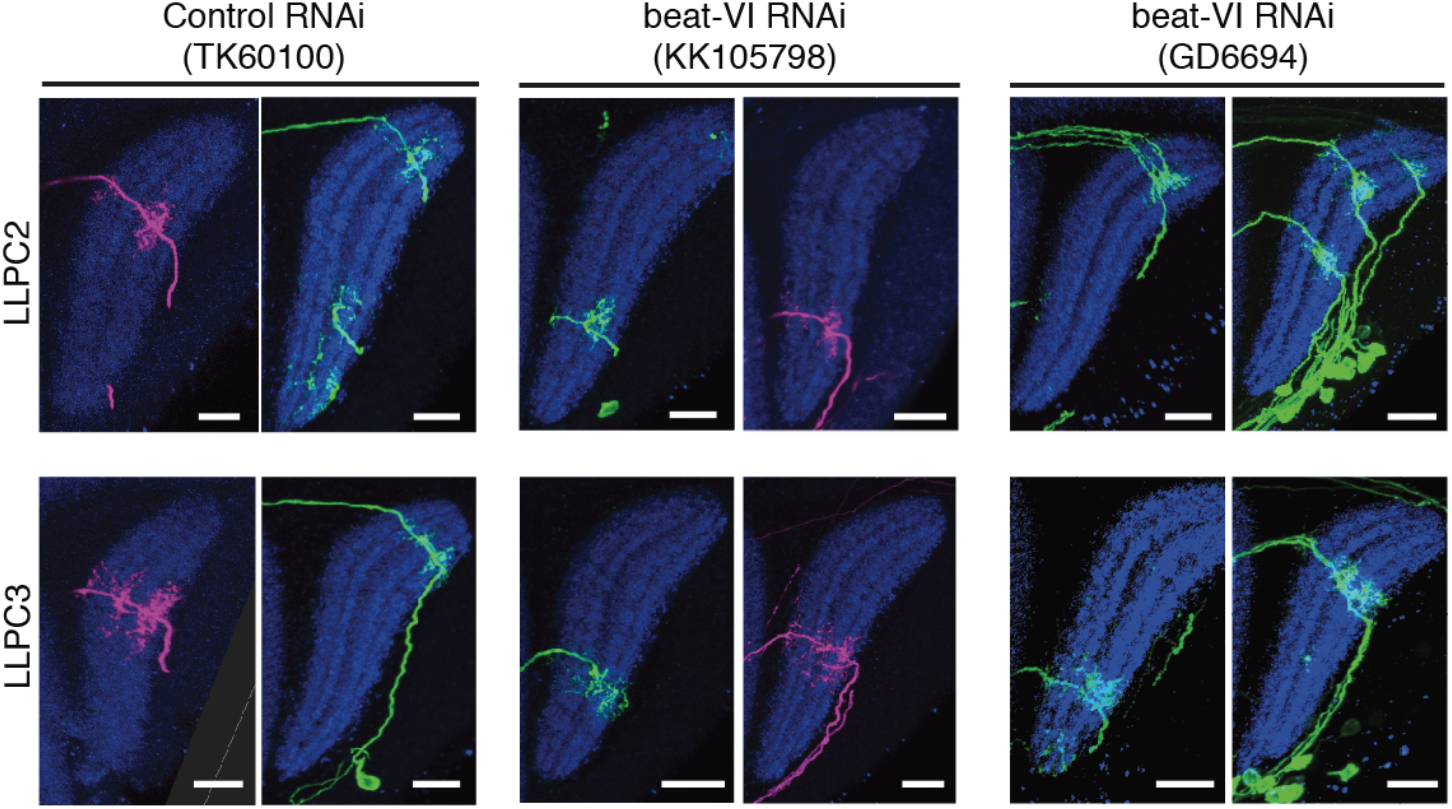
LLPC2 and LLPC3 beat-VI MCFO-RNAi. Additional examples of sparsely labeled LLPC2 and LLPC3 neurons coupled with beat-VI RNAi (same experiment as in Figures 4E-4F). Data is shown for two independent beat-VI RNAi (KK105798 is used in Figure 4). Neuropil marker (blue), brp. Scale bars, 10 μm.

**Figure S6.**
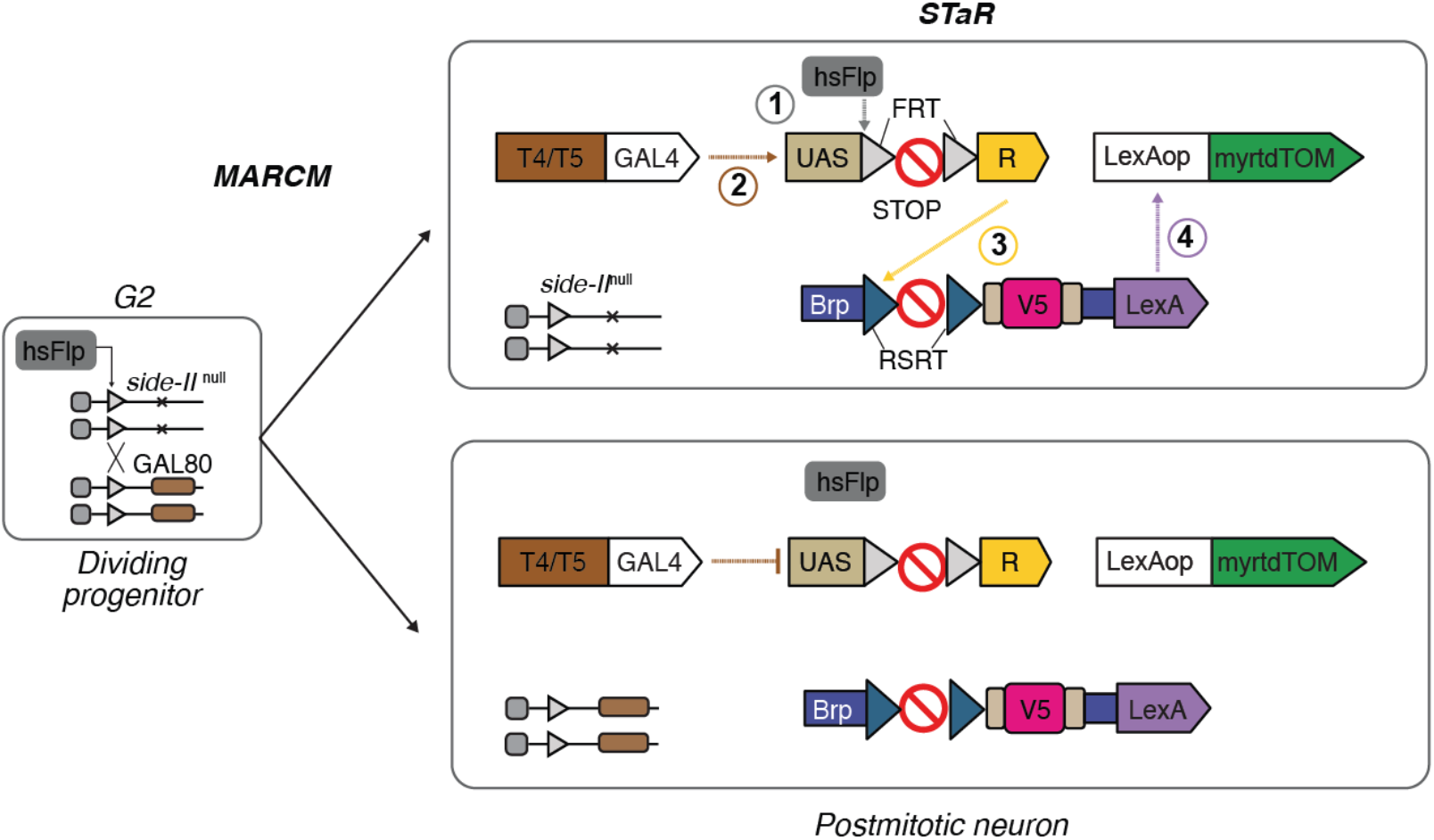
The MARCM-STaR methods. MARCM-STaR labels cell morphology and the presynaptic active zone protein Brp in single sparsely distributed null mutant neurons in otherwise wild-type backgrounds. Heatshock Flp induces mitotic recombination (MARCM) to generate homozygous null mutant clones. In the post mitotic mutant neuron where T4/T5-GAL4 is active, heatshock Flp excises the stop signal by mediating recombination between the flanking FRT recombination sites. This allows for expression of R recombinase. R recombinase, in turn, excises the stop cassette in the Brp gene by mediating recombination between RSRT sites. This recombination event inserts the V5 epitope tag into Brp and through T2A sequence (not shown) allows for co-expression of LexA; LexA then drives expression of myrtdTOM.

